# The anti-mycobacterial activity of *Artemisia annua* L is based on deoxyartemisinin and artemisinic acid

**DOI:** 10.1101/2020.10.23.352500

**Authors:** Sumana Bhowmick, Rafael Baptista, David Fazakerley, Kezia E. Whatley, Karl F. Hoffmann, Jianying Shen, Luis A. J. Mur

## Abstract

The discovery of antimalarial artemisinin from *Artemisia annua* L. is an example of how Traditional Chinese Medicine (TCM) may be exploited to meet a recognized need. In this study, we systemically investigated *A. annua* L. for its antimicrobial activity and assessed it as a source of bioactive natural products for anti-mycobacterial activity.

We used a silica gel column to perform antimicrobial activity-guided purification of the *A. annua* leaf, whose identity was confirmed by *rbcL* DNA barcoding, and used UHPLC-HRMS and NMR to elucidate the structure of purified active compounds. The antimicrobial activity of crude extracts, isolated compounds and the control artemisinin (Apollo Scientific Ltd) was assessed against *Escherichia coli*, *Pseudomonas aeruginosa*, *Staphylococcus aureus*, methicillin-resistant *Staphylococcus aureus* (MRSA), *Mycobacterium smegmatis* strains by serial micro dilution method (31.25-1000 μg/mL). The isolated compounds were tested for synergistic effects against mycobacterium.

Bioactive compounds were purified and identified as deoxyartemisinin and artemisinic acid. Artemisinic acid (MIC 250 μg/mL) was more effective in comparison to deoxyartemisinin (MIC 500 μg/mL) and artemisinin (MIC 1000 μg/mL) against *M. smegmatis*. We used a molecular docking approach to investigate the interactions between selected anti-mycobacterial compounds and proteins involved in vital physiological functions in *M. tuberculosis*, namely *Mt*Pks13*, Mt*PknB*, Mt*PanK*, Mt*KasA*, Mt*InhA and *Mt*DprE1 and found artemisinic acid showed docking scores superior to the control inhibiters for *Mt*KasA, suggesting it to be a potential nick for further *in vitro* biological evaluation and anti-TB drug design.

## 1. Introduction

Currently, tuberculosis (TB) is the leading cause of death from an infectious disease, causing 10 million incident cases globally in 2018. There were an estimated 1.2 million (range, 1.1–1.3 million) TB deaths among HIV-negative people in 2018 (27% reduction from 1.7 million in 2000), and an additional 251 000 deaths (range, 223 000–281 000) among HIV positive people (a 60% reduction from 620 000 in 2000) [1]. There remains an urgent need to discover and develop new anti-TB drugs, particularly to target drug-resistant and dormant strains of *M. tuberculosis* as well as providing a more effective and shorter duration of treatment [2].

Traditional Chinese Medicine (TCM) is one of the most comprehensive, well-documented traditional and folk medicines in human history, representing an important corpus of “folk-medicine” than could be seen as the product of centuries of “trial-and-error” approaches used by doctors. Therefore it is attracting considerable interest throughout the world. With the increased export of Chinese medicines, it has been easier to apply the modern research ways to detect the molecules, define their mode of action and pharmacokinetic and pharmacodynamic properties, and study the natural products in Chinese herbs as one of the most important resources for developing new lead compounds.

*Artemisia annua* L., a Chinese herb, is known to have anti-malarial, antipyretic, analgesic and anti-inflammatory activities. The most potent anti-malarial natural product artemisinin was purified from *A. annua* L. Artemisinin and its semi-synthetic derivatives are now front line anti-malarial drugs. A common structural feature of artemisinin derivatives with antimalarial activity is the endoperoxide bridge [3]. The antimalarial mechanism of these artemisinin derivatives appears to be dependent on the breaking of an endoperoxide bridge due to haem-mediated free-radical generation [4]. Artemisinin also exhibits activity against other protozoan parasites including the blood fluke species of *Schistosoma* [5]. The antimicrobial activity of artemisinin derivatives remains poorly defined but conjugation to a siderophore improves its anti-tubercular activity [6].

Given the wide-ranging activities of artemisinins [7] and anti-mycobacterial properties of conjugated artemisinin, we sought to identify artemisinin-like molecules from *A. annua* that, by themselves, contain potent anti-microbial activity. Thus, we isolated a small library of compounds with anti-mycobacterial activity. Two compounds were identified as deoxyartemisinin and artemisinic acid , which whilst have some structural similarity to artemisinin, lack the endoperoxide bridge.

## 2. Results

### 2.1. Antimicrobial Assay

Based on antimicrobial activity -guided purification (Supplementary Figure 1), two weakly active components were isolated and purified from crude extracts of dried A. annua leaves, and identified as deoxyartemisinin and artemisinic acid (Figure. 1, Figure 2 and Figure 3). The antimicrobial properties of deoxyartemisinin and artemisinic acid were assessed against our panel of bacterial strains (Table 1), and showed that artemisinic acid and deoxyartemisinin had MICs of 250 and 500 μg/mL against *S. aureus*. Also, both deoxyartemisinin and artemisinic acid exhibited a MIC of 500 μg/mL against *M. smegmatis* (NCTC 333) and 500 and 250 μg/mL against *M. smegmatis* (mc^2^155) respectively. However, crucially, this MIC decreased 4-fold in assessments for synergy between deoxyartemisinin and artemisinic acid. The checkerboard assay, against *M. smegmatis* mc^2^155, proved a synergistic effect between deoxyartemisinin and artemisinic acid with FICI = 0.5 and the MIC was reduced by a factor of 4 to 125 μg/mL.

**Figure 1:**
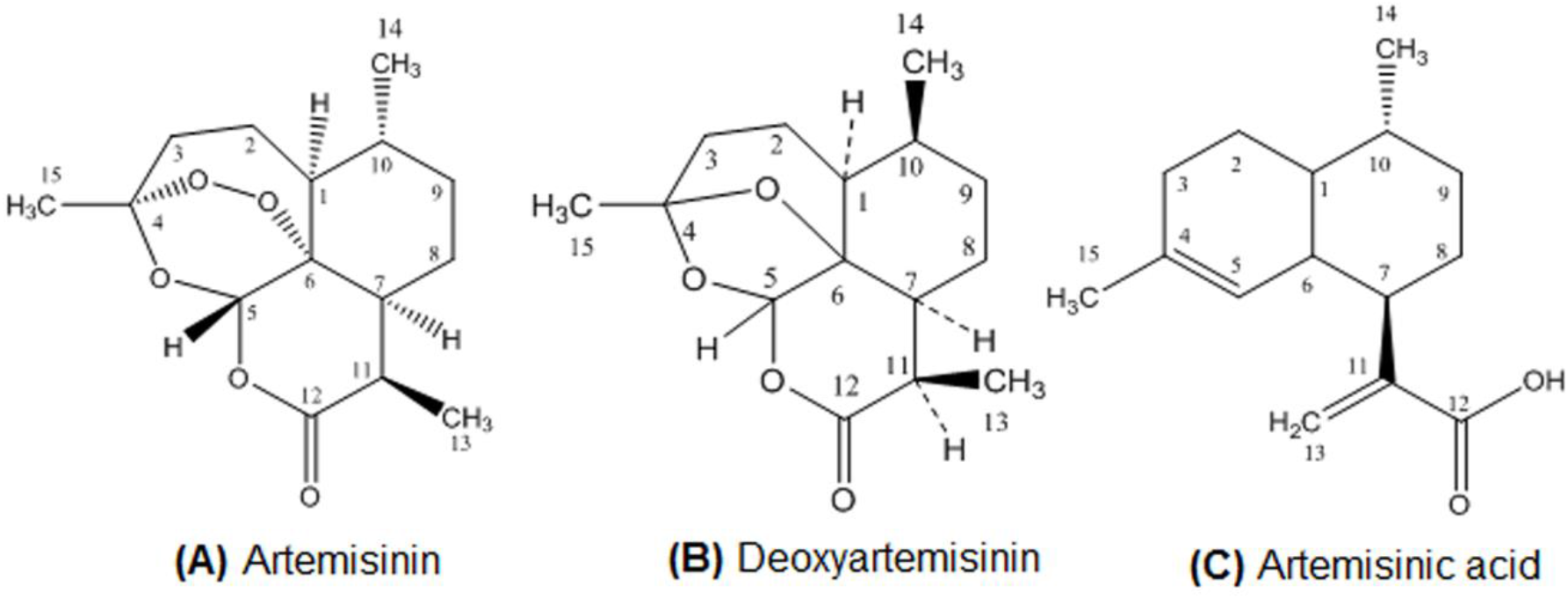
The chemical structures of three bioactive natural products isolated from *Artemisa annua.*

**Figure 2:**
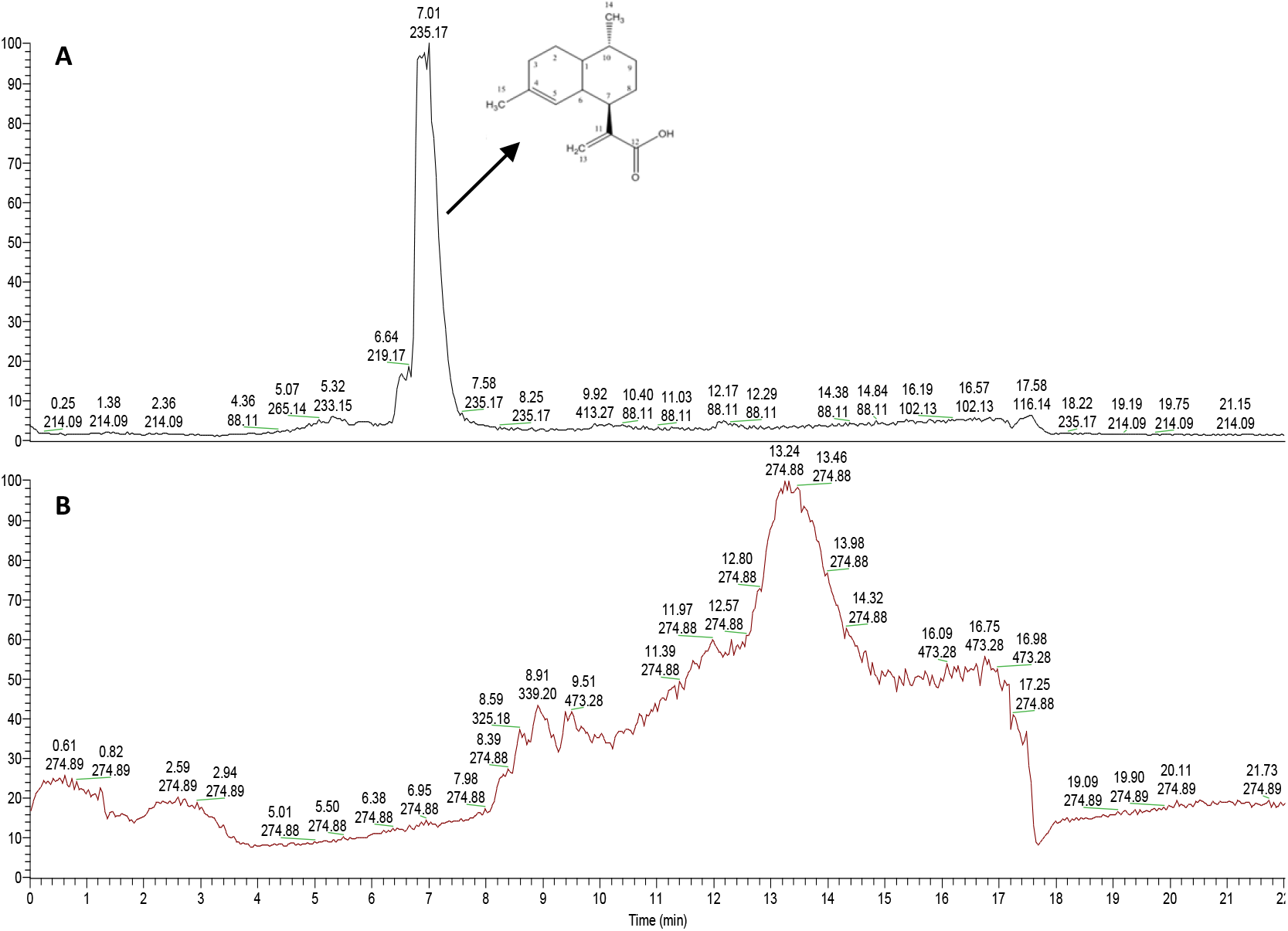
Total ion chromatograms of HFE3f. A: Positive ionization, B: Negative ionization

**Figure 3:**
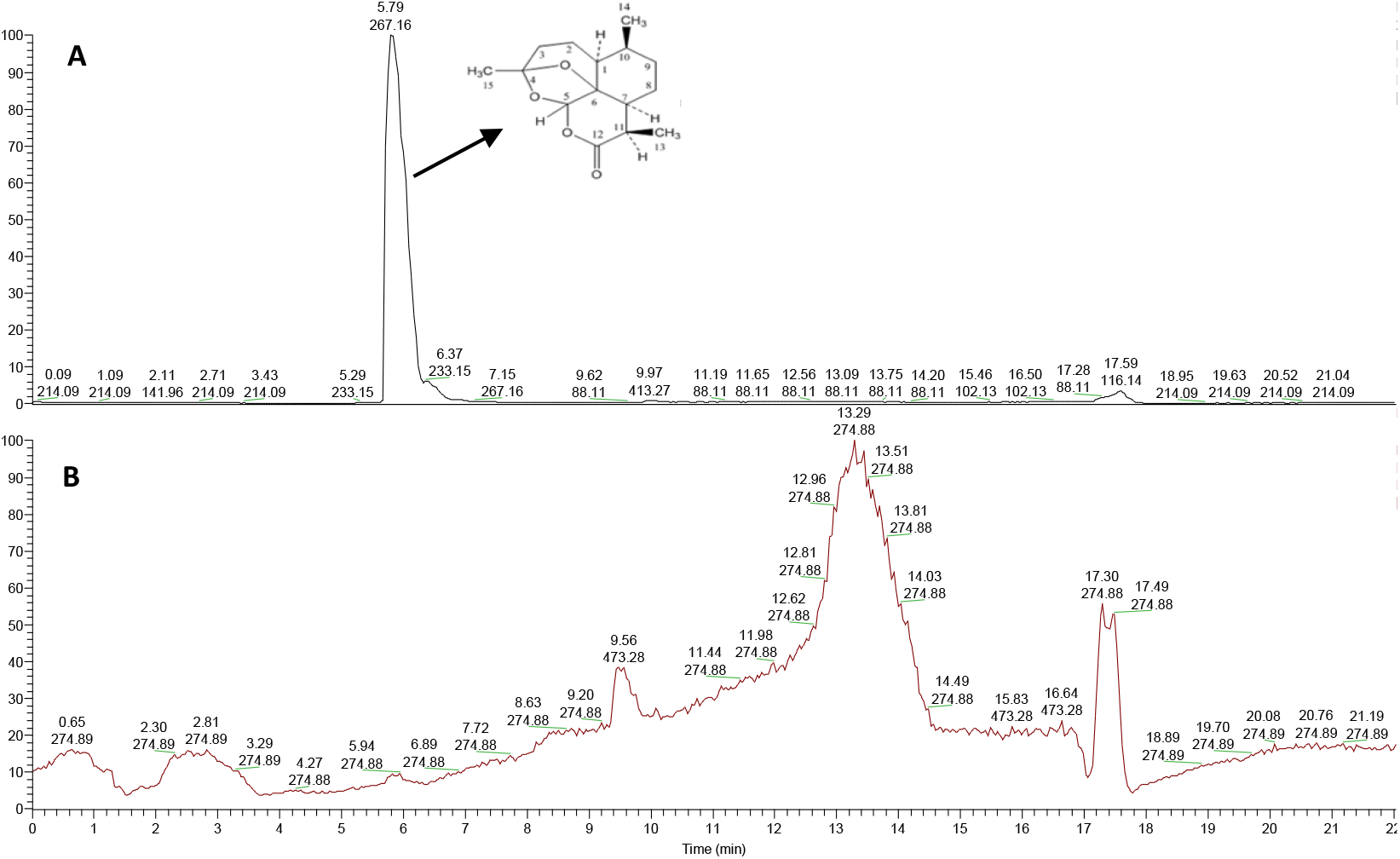
Total ion chromatograms of HFE3e. A: Positive ionization, B: Negative ionization

**Table 1:** Antibacterial activity (MIC μg/mL) of compounds 1 (deoxyartemisinin), 2 (artemisinic acid) against Gram-positive and Gram-negative bacteria

|  | Deoxyartemisinin | Artemisinic acid | Artemisinin | Rifampicin | Gentamycin Sulphate |
| --- | --- | --- | --- | --- | --- |
| *Escherichia coli* | 500 | $\geq 1000$ | $\geq 1000$ | NT<sup>1</sup> | 1.6 |
| *Pseudomonas aeruginosa* | 500 | $\geq 1000$ | $\geq 1000$ | NT<sup>1</sup> | 3.13 |
| *Staphylococcus aureus* | 500 | 250 | $\geq 1000$ | NT<sup>1</sup> | 3.13 |
| *Mycobacterium smegmatis* | 500 | 500 | $\geq 1000$ | 1.14 | NT<sup>1</sup> |
| *Mycobacterium smegmatis* mc<sup>2</sup>155 | 500 | 250 | 500 | 1.41 | NT<sup>1</sup> |
| Methicillin-resistant *Staphylococcus aureus* (MRSA) | 500 | 250 | $\geq 1000$ | NT<sup>1</sup> | 3.13 |

<sup>1</sup> NT: not tested

### 2.2. Computational studies

To suggest a mode of action, the individual binding energies of artemisinin, deoxyartemisinin and arteminisic acid were analysed by docking against *Mt*DprE1, *Mt*InhA, *Mt*KasA, *Mt*PanK, *Mt*PknB and *Mt*Pks13. These were compared with their respective positive controls which were the established inhibitor (control) drugs for each mycobacterial target. AutoDock Vina was used to predict binding affinities. Individual binding energy of artemisinin, artemisinic acid and deoxyartemisinin was analysed by docking against *Mt*DprE1, *Mt*InhA, *Mt*KasA, *Mt*PanK, *Mt*PknB and *Mt*Pks13, and compared with the respective control inhibitor, retrieved from Protein Data Bank (PDB). AutoDock Vina was used to predict binding affinities (Table **2**).

**Table 2:** Binding energies (kcal.mol^−1^) of compounds and their respective controls

| Targ<br>ets | Artem<br>isinic<br>acid | Artemi<br>sinin | Deoxy-Art<br>emisinin | Respective controls |
| --- | --- | --- | --- | --- |
| MtPks13 | -6.7 | -7.6 | -8.2 | I28 (ethyl 5-hydroxy-4-[(4-methylpiperidin-1-yl)methyl]-2-phenyl-1-benzofuran-3-carboxylate): -10.5 |
| MtPknB | -6.7 | -8.4 | -8 | Mitoxantrone: -7.7 |
| MtPanK | -7.6 | -8.7 | -8.7 | ZVT (2-chloro-N-[1-(5-[[2-(4-fluorophenoxy)ethyl]sulfanyl]-4-methyl-4H-1,2,4-triazol-3-yl)ethyl]benzamide): -8.3 |
| MtKasA | -8.3 | -6.4 | -6.9 | Thiolactomycin: -6.57 |
| MtInhA | -8 | -8.9 | -8.5 | Isoniazid: -10.4 |
| MtDprE1 | -7.2 | -8.3 | -8.3 | Bedaquiline: -10.1 |

Comparing the MIC data artemisinic acid shows lower binding energy against *Mt*KasA in comparision to artemisinin and deoxyartemisinin and also the respective control thiolactomycin (Table **2**). To further investigate the key properties necessary for an optimal binding, the chemical space spanned by the compounds was studied for all the compounds. The physiochemical properties were studied using PaDel-Descriptor including: molecular weight (MW), partition coefficient (xLogP), rotatable bonds (nRotB), H-bond donors (nHBDon_Lipinski), H-bond acceptors (nHBAcc_Lipinski) and topological polar surface area (TopoPSA) (Supplementary Figure 2).

After analysing the binding energies and physiochemical properties, a structural study was carried out based on their binding energy by docking the compounds onto *Mt*KasA and their interaction were further investigated (Figure 4) (Supplementary Figure 4).

**Figure 4:**
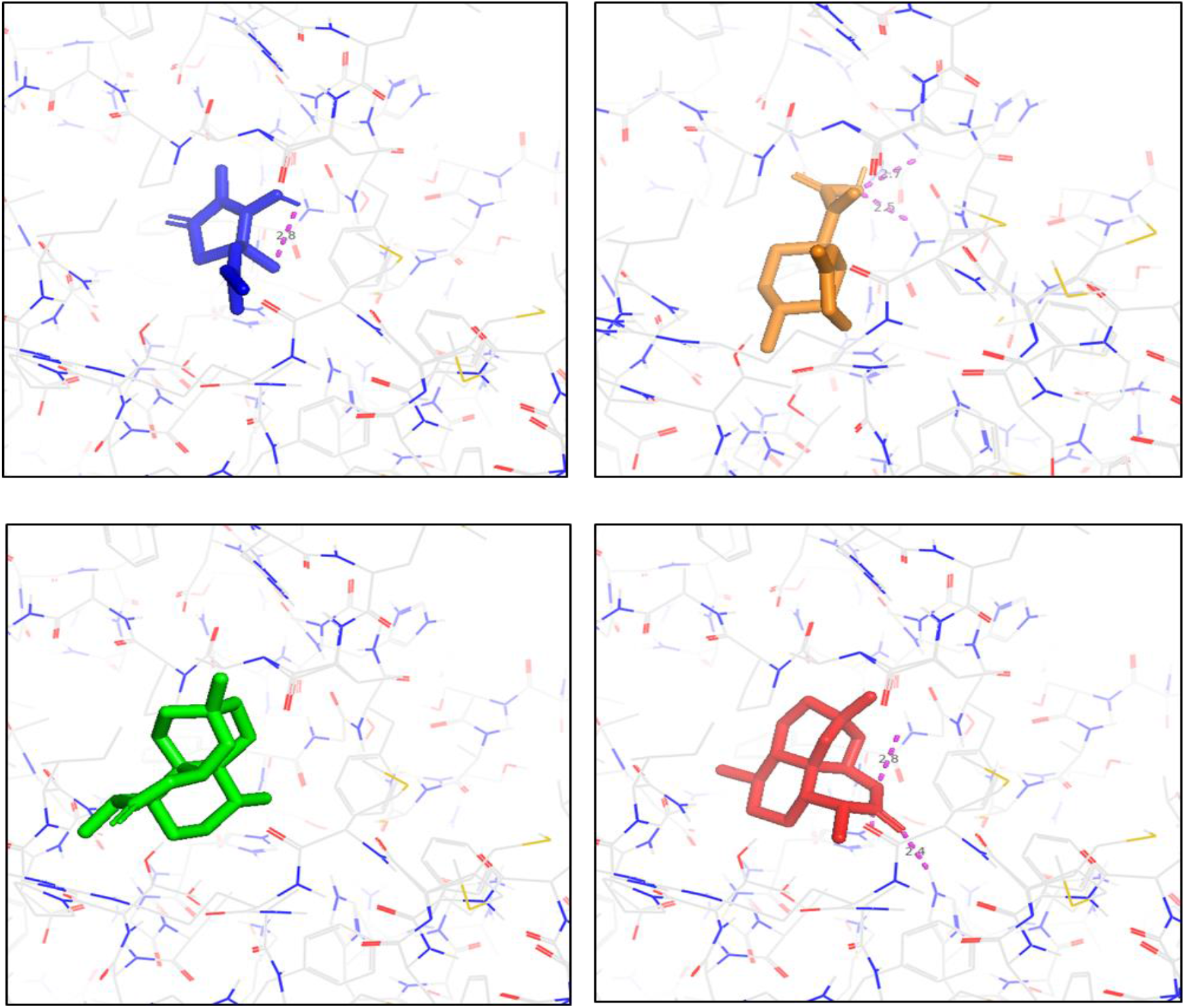
Molecular interactions of the best docking positions of Thiolactomycin (blue: top left), Artemisininc acid (Orange: top right) Artemisinin (green: down left) and Deoxyartemisinin (down right).

A closer look at the interactions between artemisinic acid and *Mt*KasA reveals that the C-12 carboxilic acid end of the molecule have a strong hydrogen bond with nitrogen (contact distance 3.0 Å) and oxygen (contact distance 2.77 Å) of Gly403 and oxygen (contact distance 3.25 Å) Asn 408. While the 4,7-dimethyl-1,2,3,4,4a,5,6,8a-octahydronaphthalen-1-yl group binds to the hydrophobic pocket of Pro280, Phe 402 and His311(Figure 5). The active site, and amino acid are in close similarity to what has been previously described as the mode of binding of the TLM control [8]

**Figure 5:**
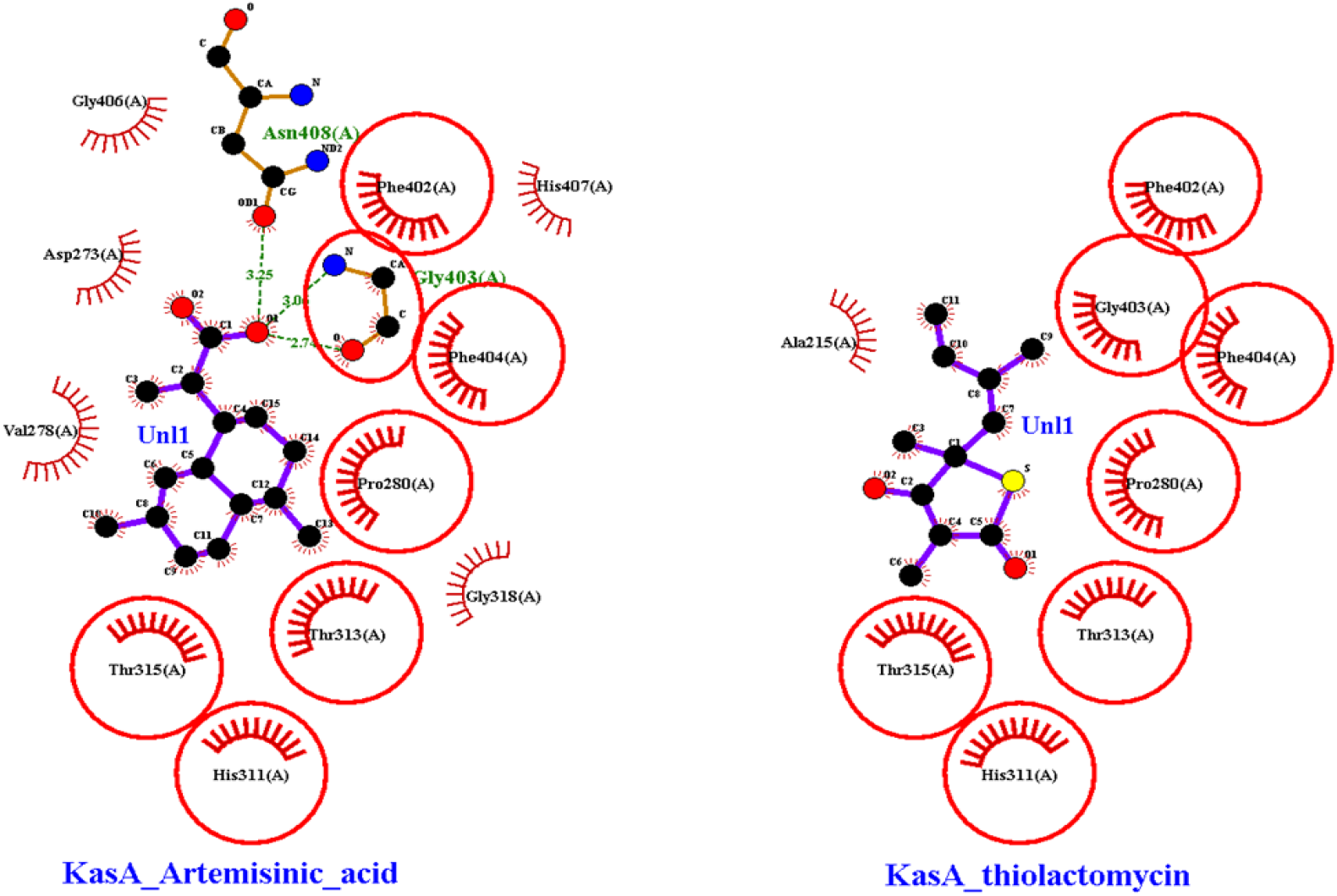
Ligplot illustration of artemisinic acid and thiolactomycin with *Mt*KasA. H-bonds in green dash. Carbons are in black, nitrogens in blue and oxygens in red. Red spheres are hydrophobic interactions

### 2.3. Figures, Tables and Schemes

## 3. Discussion

Artemisinin, deoxyartemisinin and artemisinic acid were originated from *A. annua* L.. Artemisinin and its several derivatives are commonly used to treat malaria, primarily as part of drug combination therapies. In antimicrobial studies against *S. aureus*, deoxyartemisinin and artemisinin have been reported with a MIC of 1000 and 2000 μg/mL respectively [9]. These MICs were higher than those reported for these compounds against other bacterial species [9]. Such results could reflect the strains that we used and also the penetration of compounds through the microorganism membrane, as we can clearly observe lower MICs with *M. smegmatis* mc^2^155. However, crucially, this MIC decreased 4-fold in assessments for synergy between deoxyartemisinin and artemisinic acid. The checkerboard assay, against *M. smegmatis* mc^2^155, proved a synergistic effect between deoxyartemisinin and artemisinic acid with FICI = 0.5 and the MIC was reduced by a factor of 4 to 125 μg/mL. These antimicrobial activities were significantly better than shown by artemisinin, including *M. smegmatis*. No anti-mycobacterial activity has previously been demonstrated for these natural products. We observed that the anti-mycobacterial activity of both Artemisia-derived natural products was not dependent on an endoperoxide bridge. Artemisinin, artesunate, artemisone, dihydroartemisinin and artemether, all with endoperoxide bridges, have shown to be effective against *S. mansoni*, through the peroxidative activity [10,11]. However, we found neither deoxyartemisinin nor artemisinic acid exhibited any effect on *S. mansoni* viability or mobility (data not shown), which could be caused by the lack of endoperoxide bridges in both compounds.

Furthermore, we investigated the anti TB potential of the compounds using a guided molecular docking approach to find a potential target. We selected some key enzymes required for *M. tuberculosis* to grow and survive within the eukaryotic host, that were involved in essential mycobacterial pathways and were absent from mammalian cells. The purpose of molecular docking is to use scoring algorithms to estimate the likelihood of the compound to bind to the protein ligand. Based on our MIC results and binding energies, it is likely that the artemisinic acid displayed a strong predicted binding towards with the *Mt*KasA enzyme in comparison to artemisinin, deoxyartemisinin and its control thiolactomycin. Figure 4 indicates that all the compounds accept artemisinin enter the binding site of thiolactomycin and artemisinic acid forms stronger hydrogen bonding in comparison to deoxyartemisinic acid and thiolactomycin. The docking studies provided strong evidence that the molecular basis for this activity is probably due to KasA inhibiton. However, this finding needs to be further investigated and validated *invitro* via genomics and metabolomics; followed by transformation studies.

## 4. Materials and Methods

### 4.1. Plant materials

The dried leaf material of *A. annua* was provided by Artemisinin Research Center, Institute of Chinese Materia Medica, China Academy of Chinese Medical Sciences, Beijing, China in October 2016 and was identified under light microscope by Xirong He at the Institute of Chinese Materia Medica and deposited as a voucher in that institute (Voucher Number, CMM-AberUK1). The plant sample was identified using DNA barcoding.

### 4.2. Flash column chromatography

Purification of compounds by flash chromatography was performed using a Biotage Flash chromatography system with SNAP C18 silica cartridges (30 g; 40-63 μm, 60 Å) and fraction collection controlled by photo-diode array (PDA) at 250 nm.

### 4.3. Semi-preparative high performance liquid chromatography (Semi-preparative HPLC)

Chromatographic separation was performed on a reverse phase (RP) ACE 10 C18 250×21.2 mm using water with 0.1% formic acid as a mobile phase solvent A and methanol with 0.1% formic acid as a mobile phase solvent B. Each sample was injected using a flow rate of 10 mL/min. Column oven temperature was set to 40 °C. Date was acquired with a photo-diode array (PDA) at 250 nm (Dionex).

### 4.4. Ultra high performance liquid chromatography–high resolution mass spectrometry (UHPLC-HRMS)

Fractions were analyzed on an Exactive Orbitrap (Thermo Fisher Scientific) mass spectrometer, which was coupled to an Accela Ultra High Performance Liquid Chromatography (UHPLC) system (Thermo Fisher Scientific). Chromatographic separation was performed on a reverse phase (RP) Hypersil Gold C18 1.9 μm, 2.1 × 150 mm column (Thermo Scientific) using H2O using 0.1% formic acid (v/v, pH 2.74) as the mobile phase solvent A and ACN: isopropanol (10:90) with 10 mM ammonium acetate as mobile phase solvent B. Each sample (20 μL) was analysed using 0-20% gradient of B from 0.5 to 1.5 min and then to 100% in 10.5 min. After 3 min isocratic at 100% B the column was re-equilibrating with 100% A for 7 min.

### 4.5. Spectroscopy

UV-visible spectra were determined using the Unicam UV-500. NMR spectra were obtained using Bruker Ultra shield-500 NMR spectrophotometer (1H-NMR 500 MHz, 13C-NMR 100 MHz) using either MeOD as the solvent reference.

### 4.6. DNA barcoding of plant samples

Total genomic DNA of was extracted using a DNeasy Plant Mini kit (Qiagen, UK) in accordance with the manufacturer’s instructions. The *rbcL* sequences where amplified with *rbcL*a-F: 5’-ATGTCACCACAAACAGAGACTAAAGC-3’ and *rbcL*a-R: 5’-GTAAAATCAAGTCCACCRCG-3’; DNA barcoding primers [12]. PCR used a total volume of 20 μL with 10 μL BioMix (BioLine, UK) 1 μL forward primer (10 μM), 1 μL reverse primer (10 μM), and 1 μL of the DNA template (50 ng/ μL) and 7 μL distilled water. PCR involved 1 cycle (94°C for 3 min), 35 cycles (94°C for 1 min, 55°C for 1 min, and 72°C for 1 min), and 1 cycle 72°C for 7 min. The resulting 600 bp band was sequenced in both directions on an ABI3730X, using the same primers as used for PCR. The derived sequence was submitted to BLAST to confirm 100% identity to *rbcL* sequences from *A. annua* vouchers. The *rbcL* sequence of CMM-AberUK1 has been submitted to Genbank (Accession Number MH051919).

### 4.7. Extraction and Isolation

Dried leaf material of *A. annua* was extracted sequentially using *n-*hexane, dichloromethane (DCM), and methanol (MeOH) at room temperature with continuous stirring for 24 h respectively. Bioactivity-linked fractionation of the n-hexane extract was based on the ability to suppress the growth of *M. smegmatis.*

The *n-*hexane extract was fractionated on a silica gel column chromatography (CC) (4×50 cm, 150 g of SiO2), eluted with *n-*hexane-EtOAc (1:0 to 0:1, at 5% gradient, 300 mL of each eluent), and EtOAc-MeOH (1:0 to 8:1, 300 mL of each eluent) to yield 14 fractions (designated H for “*n-*hexane” and A through to N; thus HA through to HN). The fraction MICs indicated that HF displayed the highest anti-mycobacterial activity. Fraction HF was again subjected to silica CC (1.5×50 cm, 60 g of SiO2) eluting with *n-*hexane-EtOAc (1:0 to 0:1, 5% gradient, 300 mL of each eluent) and EtOAc-MeOH (1:0 to 8:1, 300 mL of each eluent) to yield 6 crude fractions (A through F; thus HFA-HFF). The anti-mycobacterial screen suggested that fraction HFE had the highest activity (Supplementary Figure 1). The HFE fraction was then using reversed-phase flash chromatography, eluted with a methanol-water gradient resulting in 7 fractions (HFE1 through HFE7. Fractions HFE3 and HFE7 had the highest anti-mycobacterial activity. However, fraction HFE7 was not further studied due to very low yields of isolated compounds. HFE3 was then subjected to further fractionation fractionation in a Semi-preparative HPLC, to yielding 8 fractions (HFE3a through HFE3h).

All the fractions after purification was run through UHPLC-MS to analyse their mass ions. Fraction HFE3e was found to be **1**(6 mg) and HFE3e was **2**(4.5 mg) at > 95% purity. The identification of the pure compounds was made based on their accurate mass and 1D-NMR comparison to previous studies [9,13].

**1**: 1H-NMR (500 MHz, CDCl3) δ 0.93 (3H, s, H-14), 0.99 (2H, m, H-8), 1.11 (1H, m, H-9), 1.19 (3H, d, J = 7.2 Hz, H-13), 1.23 (1H, m, H-2), 1.25 (1H, m, H-10), 1.27 (3H, m, H 1), 1.51 (3H, s, H-15), 1.61 (1H, m, H-3), 1.79 (2H, m, H-3’ and H-9’), 1.90 (2H, m, H-2’ and H-8’), 1.99 (1H, m, H-7), 3.17 (1H, m, H-11), 5.69 (1H, s, H-5) ppm; 13C-NMR (100 MHz, CDCl3) δ 12.74 (C-13), 18.69 (C-14), 22.20 (C-2), 23.68 (C-8), 24.09 (C-15), 32.92 (C-11), 33.66 (C-9), 34.17 (C-3), 35.53 (C-10), 42.64 (C-7), 44.85 (C-1), 82.58 (C-6), 99.80 (C-5), 109.32 (C-4), 171.88 (C-12) ppm.

**2**: 1H-NMR (500 MHz, MeOD) δ 0.92 (3H, d, J = 6.3 Hz, H-14), 1.07 (1H, m, H-9), 1.30-1.45 (2H, m, H-3), 1.40 (3H, m, H-1 and H-8), 1.55 (1H, m, H-10), 1.59 (3H, s, H-15), 1.70-1.83 (2H, m, H-2), 1.90 (1H, m, H-9’), 1.98 (1H, m, H-6), 2.68 (2H, m, H-7), 5.02 (1H, s, H-5), 5.45 (1H, s, H-13), 6.29 (1H, s, H-13’) ppm; 13C-NMR (100 MHz, MeOD) δ 20.2 (C-14), 23.8 (C-15), 26.7 (C-2), 27.1 (C-8), 27.3 (C-3), 28.9 (C-10), 36.5 (C-9), 39.4 (C-6), 43.1 (C-1), 43.7 (C-7), 121.5 (C-5), 124.6 (C-13), 135.9 (C-4), 145.3 (C-11), 170.7 (C-12) ppm.

### 4.8. Antimicrobial assays

The antimicrobial activity of all the extracts and isolated compounds were assessed against *Escherichia coli* ATCC 25922, *Pseudomonas aeruginosa* ATCC 27853, *Staphylococcus aureus* ATCC 29213, methicillin-resistant *S. aureus* (MRSA), *Mycobacterium smegmatis* strains NCTC 333 and mc^2^155. Rifampicin and gentamycin sulphate were the reference antimicrobial. The checkerboard method and the fractional inhibitory concentration index (FICI) was used to determine potential synergistic, additive or even antagonistic effects of combinations of individual compounds at different concentrations as previously described [14].

### 4.9. In silico studies

All chemical structures were retrieved from the PubChem compound database (NCBI) (https://www.pubchem.ncbi.nlm.nih.gov). The crystal structures and respective controls of MtDprE1 (PDB ID: 6HEZ)[15], MtInhA (PDB ID: 1ENY) [16], MtKasA (PDB ID: 2WGE) [17], MtPanK type 1 (PDB ID: 4BFT) [18], MtPknB (PDB ID: 2FUM) [19] and MtPks13 (PDB ID: 5V3X) [20] were retrieved from the RCSB Protein Data Bank (PDB) database (https://www.rcsb.org).

*In silico* prediction of physicochemical and structural properties of the compounds was performed using PaDEL-Descriptor [21] including the descriptors: nHBAcc_Lipinski (acceptor H-bonds), nHBDon_Lipinski (donor H-bonds), nRotB (number of rotation bonds), TopoPSA (topological polar surface area), MW (molecular weight) and XLogP (prediction of logP based on the atom-type method).

An extended PDB format, termed as a PDBQT file, was used for coordinate files which includes atomic partial charges [22]. All file conversions were performed using the open source chemical toolbox Open Babel 2.3.2 [23]. The ligand and protein structures were optimised using AutoDock Tools software (AutoDock 1.5.6) which involved adding all hydrogen atoms to the macromolecule, which is a step necessary for correct calculation of partial atomic charges. Gasteiger charges are calculated for each atom of the macromolecule in AutoDock 1.5.6 [22].

All the compounds were docked against *Mt*DprE1, *Mt*InhA, *Mt*KasA, *Mt*PanK, *Mt*PknB,and *Mt*Pks13 respectively along with the controls. Molecular docking calculations for all compounds with each of the proteins were performed using AutoDock Vina 1.1.2. Docking calculation was generated with the software free energy binding own scoring function. The binding affinity of the ligand was observed as a negative score with units expressed as kcal.mol^−1^. Nine different poses were calculated for each protein with the parameters num_modes = 9 and exhaustiveness = 16. The lowest energy conformation was chosen for binding model analysis. Molecular interactions between ligand and protein were generated, analysed by LigPlot+ and depicted by PyMOL. PyMOL Molecular Graphics System, Version 2.0 Schrödinger (http://www.pymol.org) was used to prepare figures [24].

To provide enough space for free movements of the ligands, the grid box was constructed to cover the active sites and was defined using AutoDock 1.5.6. The grid points for *Mt*DprE1, the grid points were set to 20 × 20 × 20, at a grid center of (x,y,z) 14.99, −20.507, 37.226 with spacing of 1 Å. For *Mt*InhA, the grid points were set to 26 × 24 × 22, at a grid center of (x,y,z) −5.111, 33.222, 13.410 with spacing of 1 Å. For *Mt*KasA, the grid points were set to 20 × 20 × 20, at a grid center of (x,y,z) 38.342, −7.033, 13.410 with spacing of 1 Å. For *Mt*PanK, the grid points were set to 20 × 20 × 20, at a grid center of (x,y,z) −18.742, 13.919, 11.679 with spacing of 1 Å. For *Mt*PknB, the grid points were set to 21 × 20 × 20, at a grid center of (x,y,z) 61.518, 2.429, −25.588 with spacing of 1 Å. For *Mt*Pks13 the grid points were set to 16 × 18 × 14, at a grid center of (x,y,z) 3.954, 27.324, 8.499 with spacing of 1 Å.

## 5. Conclusions

Our studies have demonstrated that the anti-mycobacterial activity of *Artemisia annua* L is based on deoxyartemisinin and artemisinic acid, but not artemisinin. It clearly indicated endoperoxide bridge-independent activity that was exhibited at least against Mycobacterium. The interation of KasA and the compounds is to be verified based on efficacy; toxicity and pharmacokinetic properties is needed be validated experimentally. It is possible that these natural products or their target could be exploited in the search for new anti-TB drugs.

## Supplementary Materials

The following are available online at www.mdpi.com/xxx/s1.

Figure S 1: Bioassay guided purification of *A.annua*

Figure S 2: Screening for anthelminthic activity against *Schistosoma mansoni* schistosomula. Using the Roboworm platform, anthelminthic activity of deoxyartemisinin (C1) and arteminisic acid (C2) were compared to those of Auranofin (AUR, 10 μM), Praziquantel (PZQ, 10μM) and the negative control dimethylsulphoxide (DMSO). (A) Images of schistosomula at 72 h following treatments. Schistosomula treated with AUR and PZQ exhibited obvious phenotypic changes compared to DMSO. Treatments (two-fold dilutions; 10 μM-0.625 μM) with C1 and C2 showed no obvious effect. Viability for (B) C1 and (C) C2 treatments as well as assessments of motility for (D) C1 and (E) C2 were quantified. All results were tested for significance using one-way ANOVA with a Dunn’s post-test. Results falling within the “hit zone” were significantly (P <0.05) from negative DMSO treated controls. In no case were significant effects for C1 and C2 observed.

Figure S 3: Physiochemical properties of Artemisinin, Artemisinic acid, Deoxyartemisinin and thiolactomycin analysed by PaDel-Descriptor.

Figure S 4: Superposition of the best docking positions of Deoxyartemisinin (Red), artemisinin(green), Artemisinic acid (orange), Thiolactomycin (Blue) against KasA

## Author Contributions

“Conceptualization, L.M. and J.S.; methodology, S.B.; R.B., K.W. resources, J.S..K.H.; writing—original draft preparation, S.B.; writing—review and editing, S.B, R.B, R.W., K.H., J.S. and L.M.; supervision, L.M. and J.S. project administration, L.M. and J.S. All authors have read and agreed to the published version of the manuscript.”

## Acknowledgments

SB is funded through an Aberystwyth University PhD Studentship and is co-supervised by LM and SJ. RB and DMF are funded through Life Sciences Research Network Wales PhD Studentships. The Roboworm platform is also funded by Life Sciences Research Network Wales. SJ’s work is supported by Fundamental Research Funds for the Central public welfare research institutes (ZZ11-046). This work was made possible through a networking Life Sciences Research Network Wales grant to LM and through BBSRC funded infrastructure at the IBERS, AU (UK). We appreciate the technical support provided by Dr. Manfred Beckmann (Aberystwyth University UK) by Prof. Robert Nash (Phytoquest, UK) and Dr. Salvatore Ferla (Cardiff University, UK).

## SUPPLEMENTARY MATERIALS

**Figure S 5:**
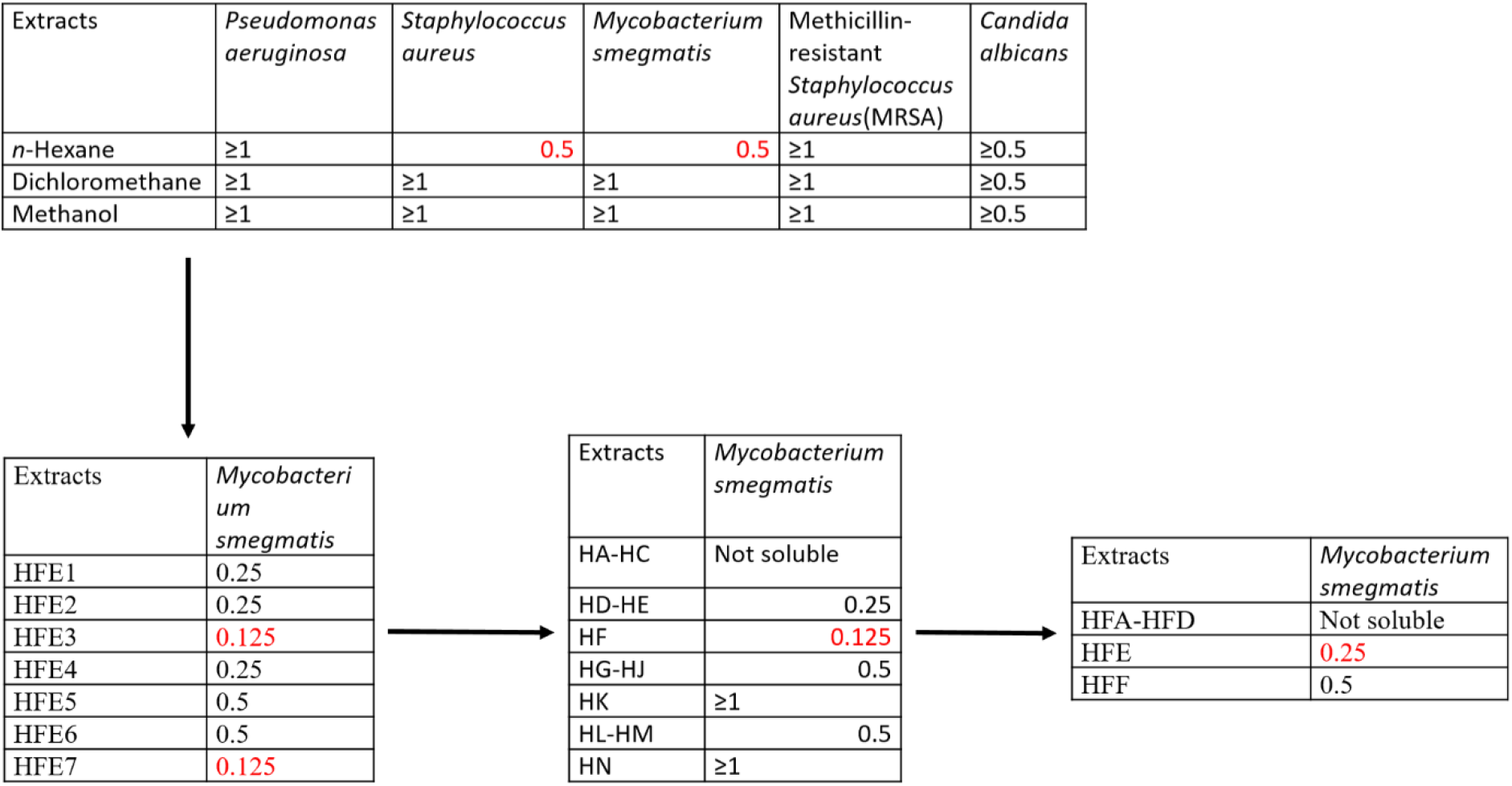
Bioassay guided purification of A.annua.

**Figure S 6:**
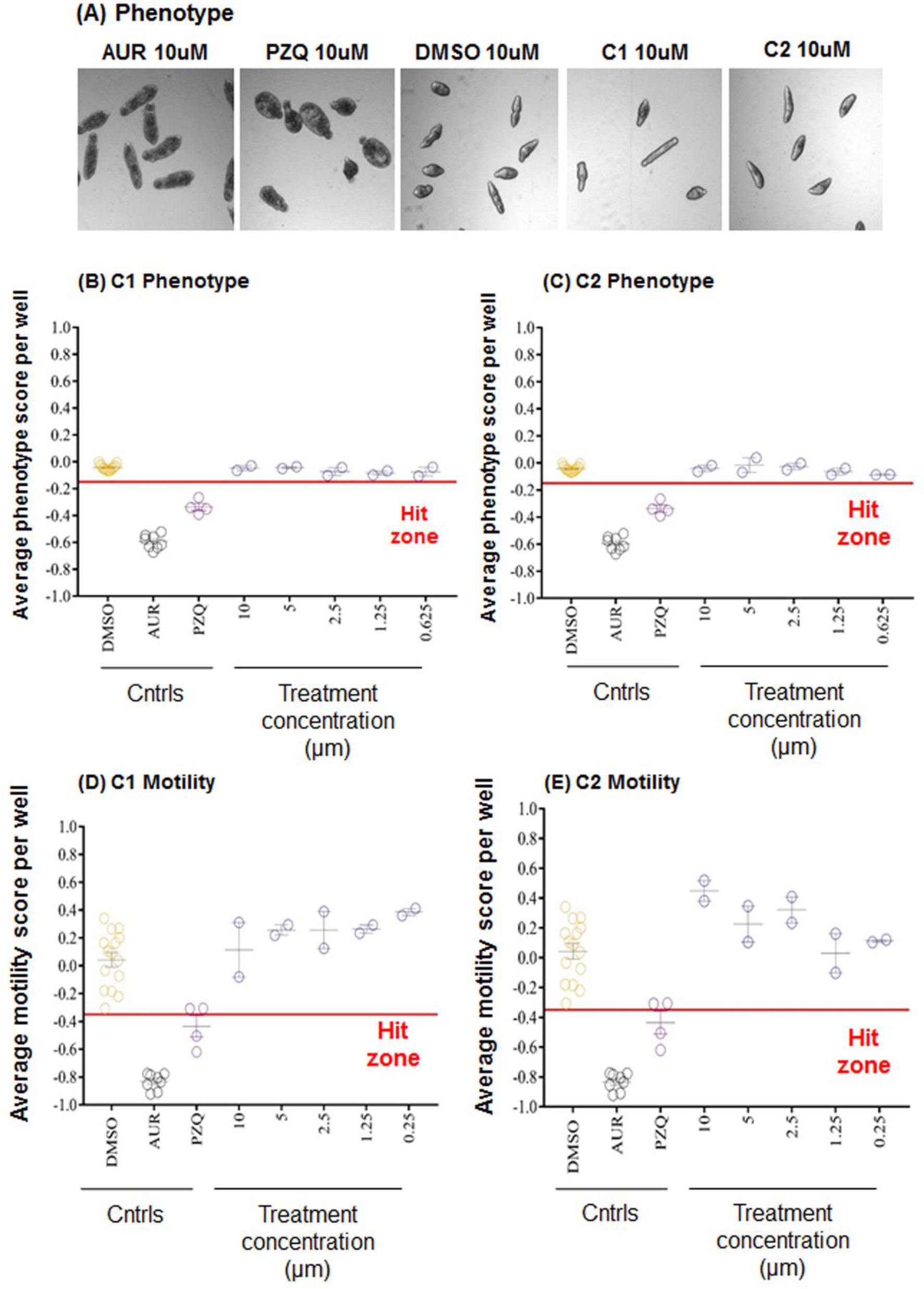
Screening for anthelminthic activity against *Schistosoma mansoni* schistosomula. Using the Roboworm platform, anthelminthic activity of deoxyartemisinin (C1) and arteminisic acid (C2) were compared to those of Auranofin (AUR, 10 μM), Praziquantel (PZQ, 10μM) and the negative control dimethylsulphoxide (DMSO). (A) Images of schistosomula at 72 h following treatments. Schistosomula treated with AUR and PZQ exhibited obvious phenotypic changes compared to DMSO. Treatments (two-fold dilutions; 10 μM-0.625 μM) with C1 and C2 showed no obvious effect. Viability for (B) C1 and (C) C2 treatments as well as assessments of motility for (D) C1 and (E) C2 were quantified. All results were tested for significance using one-way ANOVA with a Dunn’s post-test. Results falling within the “hit zone” were significantly (P <0.05) from negative DMSO treated controls. In no case were significant effects for C1 and C2 observed.

**Figure S 7:**
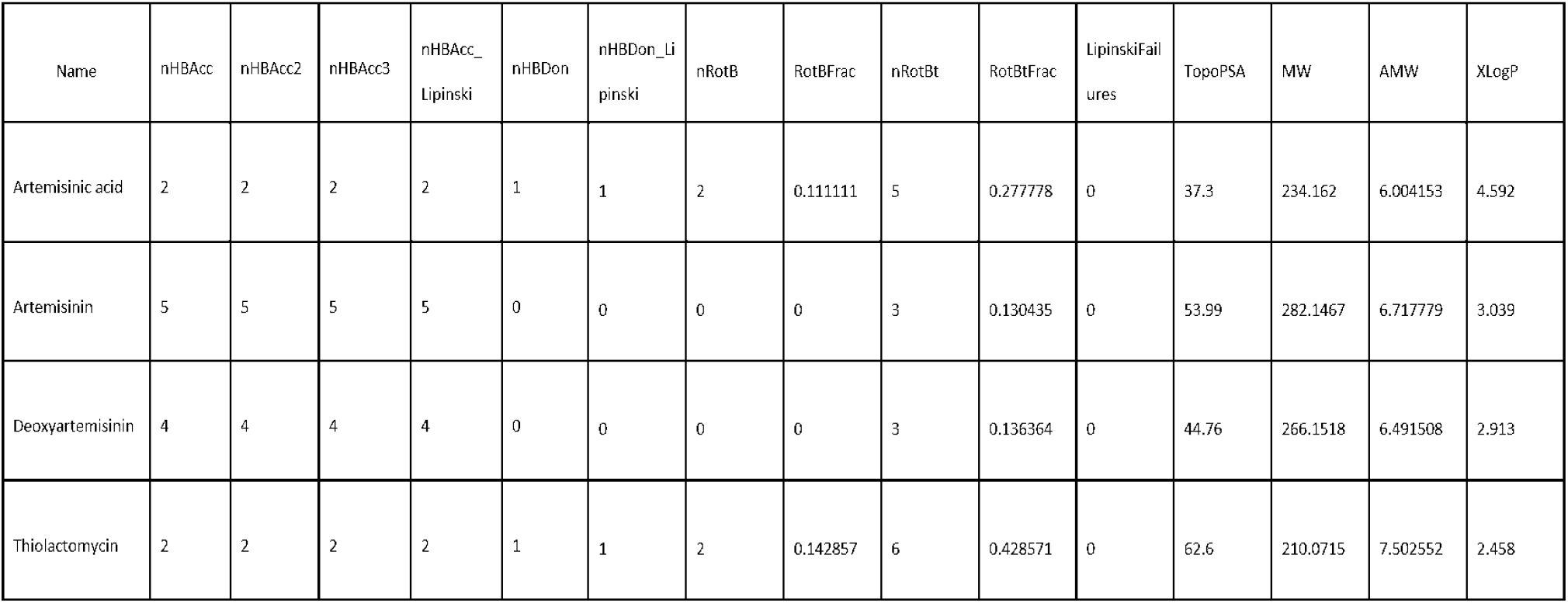
Physiochemical properties of Artemisinin, Artemisinic acid, Deoxyartemisinin and thiolactomycin analysed by PaDel-Descriptor.

**Figure S 8:**
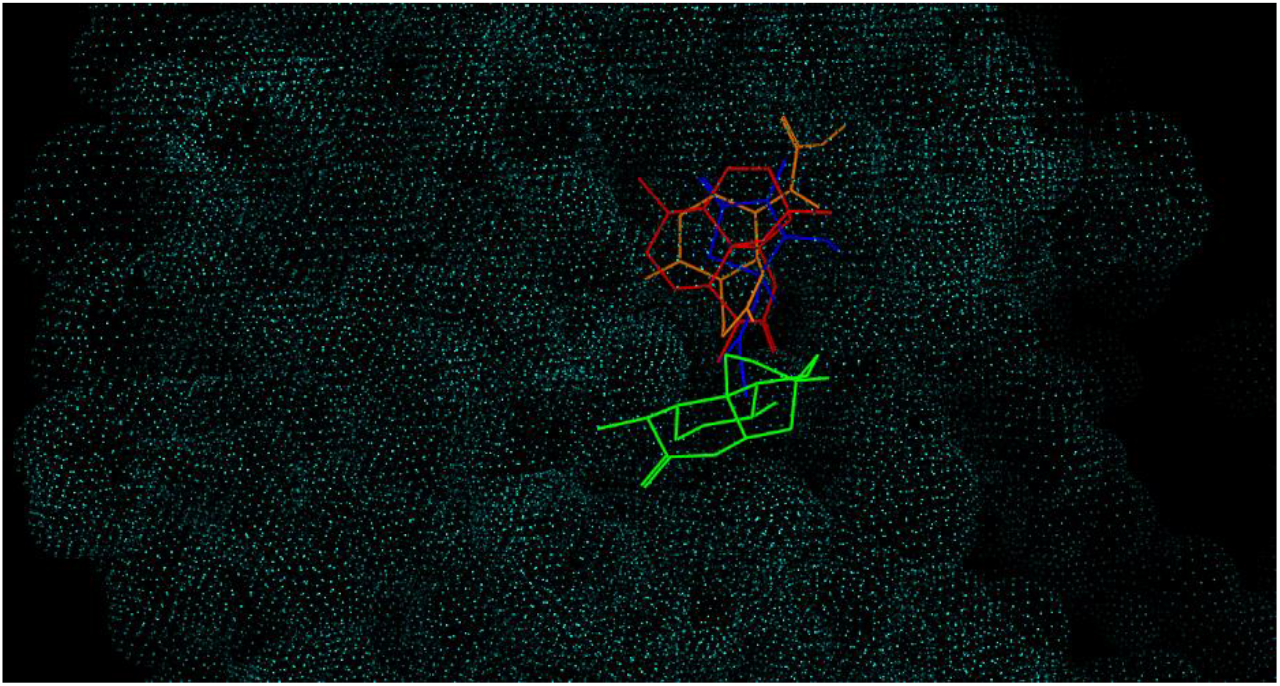
Superposition of the best docking positions of Deoxyartemisinin (Red), artemisinin(green), Artemisinic acid (orange), Thiolactomycin (Blue) against KasA opological polar surface area (TopoPSA)

**Table 3:**
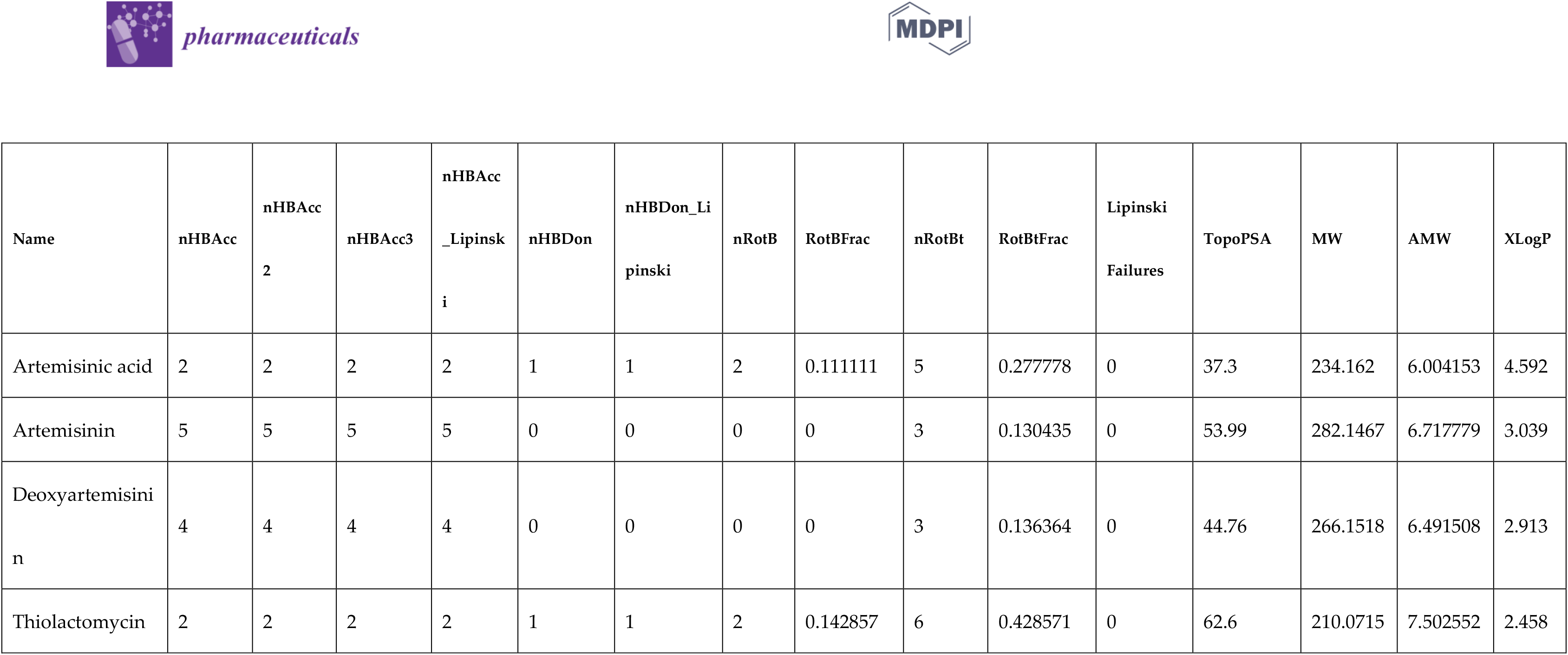
The physiochemical properties using PaDel-Descriptor: molecular weight (MW), partition coefficient (xLogP), rotatable bonds (nRotB), H-bond donors (nHBDon_Lipinski), H-bond acceptors (nHBAcc_Lipinski) and t.

